# Physiological Evaluation of Far Infrared-Emitting Garments on Sleep and Thermoregulation

**DOI:** 10.1101/2024.06.13.598953

**Authors:** Masaki Nishida, Taku Nishii, Shutaro Suyama, Sumi Youn, Kenta Funayama

## Abstract

Garments engineered to emit far-infrared (FIR) spectral frequencies have potential conditioning benefits. This study aimed to explore the physiological effects of FIR-emitting clothes on sleep propensities involving thermoregulation and autonomic nervous function. In this double-blind, randomized, placebo-controlled, crossover study, fifteen healthy male participants (age 21.6 ± 1.5 years, body mass index 23.1 ± 1.9, Pittsburgh Sleep Quality Index 5.0 ± 2.0) wore a whole-body FIR garment (wavelength: 5–20 μm at 40°C), while a total body garment with the 100% polyester fiber without ceramic components was used as control for the intervention. To compare the physiological variables between the two electroencephalograms, the tympanic membrane temperature, sweating rate, skin temperature, and humidity were measured during sleep. Heart rate variability during sleep was also calculated. The percentage of rapid eye movement (REM) sleep was significantly higher in the FIR compared than in control (*p* = 0.027). We found a significant main effect of tympanic membrane temperature in time, equivalent to core body temperature. (2-way ANOVA; (*F*(26, 234)=4.046, *p*<0.001). Furthermore, the sweating rate was significantly lower when sleeping with FIR predominantly in the final phase of sleep (garment × time interaction (*F*(26,572)=55.05, *p*<0.001). Although there were no significant differences in skin surface temperature between the two garments, FIR provided significantly lower skin humidity during the middle of night sleep compared with the control. Regarding LF, a significant garment type x time interaction (*F*(1.87, 48.57)=3.36, *p*=0.046). Post-hoc analysis revealed that the LF value of FIR was higher in the final phase of sleep than that of control (p=0.027). Then present results suggest that sleeping with FIR in comparison to with control, has a beneficial effect to facilitate restorative sleep, presumably involving proper thermoregulation.

## Introduction

Sleep is significantly affected by environment [1]. The importance of sleep environment and its beneficial physiological effects are gaining attention. Recent studies have validated the optimization of sleep environments, such as bedding, mattresses [2–4], or chairs [5]. These studies examined the usefulness of the sleep environment of bedding in terms of sleep variables and physiological indicators such as core body temperature (CBT), autonomic nervous function, and electromyography. Nevertheless, there have been few scientific studies on the optimization of clothing or garments worn during night sleep.

Sleeping garments affect several aspects of human sleep, including temperature and humidity. Changes in heat and humidity depend considerably on the materials used in sleeping garment [6]. Owing to the distinct thermal insulation and hygral characteristics inherent to each fiber type, the utilization of fabrics derived from various fiber types appears to have disparate impacts on thermal insulation [7]. Chow et al. compared sleep quality using wool, cotton, and polyester as sleeping garment materials and demonstrated that wool was superior in several sleep variables [8]. The CBT, which is essential for regulating sleep-wake rhythms, was not assessed in this study. Instead of the CBT, the body sweat evaporation rate was calculated, which showed no significant differences among the sleeping garments. Assessing current materials is challenging because of their inherent propensities, subjective preferences, and ambient conditions [9].

Various technologies related to garment materials have been developed to manipulate garments specially engineered to emit frequencies within the far-infrared (FIR) spectrum. Far infrared (FIR) is a specific band (15 to 1000 μm) in the infrared spectrum of electromagnetic radiation [10]. Any object exhibiting a temperature above absolute zero, such as the human body at room temperature, emits far-infrared FIR radiation. FIR rays can penetrate human tissues up to 4 cm, stimulating human biological functions at the cellular level [11]. FIR-emitting materials provide various biological effects on humans, noteworthily, the promotion of circulatory dynamics. A previous animal study demonstrated that FIR increase skin microcirculation involved in the biological effect of nitric oxide [12] and forearm skin blood flow [13], as well as enhance blood circulation and metabolism [11]. FIR may ameliorate sleep-wake rhythm, which predominantly determines CBT regulation by regulating vasomotor tone (dilation and constriction) [14]. Inoue et al. explored the effect of FIR on human sleep using FIR radiator disks embedded in bedclothes and showed an increase in body tissue temperature [15]. Recently, randomized controlled trials showed that FIR pyjamas had fewer beneficial effects on the subjective assessment of sleep by questionnaires, not by physiological indices [16]. Thus, the effects of FIR on human sleep propensity have not been sufficiently examined using physiological techniques.

In the current study, we explored the effects of FIR on the physiological variables associated with sleep using an FIR-emitting garment. Technology using FIR is rapidly advancing and is being applied to clothing and training wear for athletes [17].

Sleeping garments that incorporate FIR technology have been developed and are now available for daily use. This preliminary study was conducted to compare the use of a technically manufactured FIR-emitting garment sleepwear made with a control fabric during night sleep. We hypothesized that an FIR-emitting garment (1) induces proper reduction of the CBT during sleep, preferentially regulates thermoregulation, (2) improves sleep variables relating to quality, and (3) appropriately coordinates the sympathovagal balance.

## Methods & Materials

### Evaluated Materials

We tested a FIR-emitting whole-body covering garment with long sleeves and long pants marketed as BAKUNE^®^ (TENTIAL Inc. Tokyo, Japan). The FIR garments were made from special functional fiber (SELFLAME^®^), manufactured with cotton (59%), polyester (38%) and polyurethane (3%). They incorporated a micro ceramic powder which, when excited by the infrared frequency produced by the body, emits a constant infrared emission of FIR frequencies (wavelength: 5–20 μm at 40°C). A similar shape and color of garments of 100% polyester without the micro ceramic powder (P-3000^®^, wundou Corp., Tokyo, Japan) was used as the control garment. The FIR and polyester control garments were clean, plain, and visibly similar, and the product logo was concealed with a piece of taped fabric. The participants were asked to choose a garment of four sizes (small, medium, large, and extra-large) according to their body shape.

### Participants

In this study, we enrolled 15 healthy men aged 20–21 years with regular sleep schedules. Recruitment was ongoing between September 2022 to February 2023. The sample size was calculated using G*Power 3.1.9.6 statistical power analysis software [18]. Power analysis indicated that a sample size of 15 per group (number of measurements = 2) were needed for a η^2^ (0.25) when a = 0.05 for a power of 0.60 with two independent conditions, using repeated measures of variance (ANOVA), within - between interactions. Statistical significance was set at p < 0.05 (two-tailed).

Participants were recruited through poster advertisements, social media, and emails to the students at Waseda University. Participants with sleep disorders, circadian rhythm disorders, or those using medication were excluded. Participants were requested to abstain from alcohol, caffeinated beverages, and vigorous exercise during the study period. All participants were assessed using the Pittsburgh Sleep Quality Index (PSQI) for habitual sleep quality and the Epworth sleepiness score (ESS) for daytime sleepiness. For the 15 participants, the mean height was 175.1 ± 6.1 cm, weight was 71.1 ± 6.4 kg, BMI was 23.1 ± 1.9, PSQI was 5.0 ± 2.0 (range 3–9), ESS was 6.6 ± 2.6 (range 3–12).

### Procedures

The present study was a double-blind, randomized, placebo-controlled, crossover study that examined the physiological variables related to sleep under controlled conditions. During night sleep, the participants wore technical clothing made with FIR fabrics or technical clothing made with conventional polyester materials as controls.

The participants wore an MTN-221 (Acos Co., Ltd., Nagano, Japan) device on the front side of the trunk by clipping it onto their waist belt or the edge of their trousers/pants prior to the experiment to assess their average bedtime and wake time.

The recording started at 18:00 the preceding night and stopped at 08:00 on the day of the experiment. The sensitivity and specificity of MTN were equivalent to those determined for conventional actigraphy [19] and polysomnography (PSG) [20]. On the testing nights, the participants were not informed of the type of garment they would wear before sleeping. Participants ate a habitual meal approximately 4 h before their average bedtime.

Participants were asked to change their sleepwear prior to bedtime, during which time they had measuring devices attached to a quiet and dark shield room implemented for sleep monitoring in the sleep laboratory of Waseda University. The temperature and relative humidity (RH) in the shield room were maintained at ∼25 °C and 45%, respectively. Following cultural conventions, the participants wore sleepwear with cotton underwear. Participants rested on a highly resistant mattress covered by a thin cotton bedsheet and under a cotton quilt of the thickness normally used in both the FIR and control conditions. Participants completed one night of the experiment (23:00 to 7:00) for both garments at a minimum of 1-week intervals.

This study was approved by the Academic Research Ethical Review Committee of Waseda University (Approval No. 2022-037). The participants were informed in advance of the purpose, methods, and risks of the experiment, and written informed consent was obtained from them. The experimental results are presented as numbers and managed confidentially.

### Ambient Conditions

The temperature and RH levels in the shielded room were continuously monitored using a thermohygrometer (TEM-201, Tanita Corp., Tokyo, Japan). In the bedrooms, temperature and RH levels were controlled using a wall-mounted air conditioner (LC1E9, YANMAR Co., Ltd., Osaka, Japan). Before the experiment, the examinees confirmed that both the temperature and humidity could be regulated using air conditioning. Ventilation was implemented continuously, although it was adjusted such that there was no evident airflow within the shielded room. Sound stimuli from outside the room were completely blocked, and the lights were switched off at bedtime, thereby considerably dimming the monitoring room.

### Sleep Questionnaire

After waking up, the participants were asked to assess their subjective sleep quality using the Oguri-Shirakawa-Azumi sleep inventory MA version (OSA-MA), specifically the version tailored for middle-aged and elderly individuals [21]. This inventory comprises 16 self-reported items, each rated on a 4-point scale. These items were categorized into five subscales: Factor I (morning sleepiness), Factor II (initiation and maintenance of sleep), Factor III (frequency of dreams), Factor IV (feeling refreshed upon awakening), and Factor V (sleep duration). The Zi value, which is indicative of sleep quality, was computed with higher values indicating better sleep quality.

Participants completed sleep surveys using the OSA sleep questionnaire twice: before and after the intervention.

### Measurements

#### Electroencephalogram recording and sleep stage scoring

Electroencephalogram (EEG) recordings were obtained during nighttime sleep using a portable EEG recording system (Insomnograf K2; S’UIMIN Inc., Tokyo, Japan). This lightweight (162 g) recording device consisted of four EEG electrodes (Fp1, Fp2, M1, and M2) and one reference electrode (Fpz) according to the 10–20 system. To validate the sleep-staging accuracy, simultaneous recording using the portable system revealed over 95% agreement for all variables, with interclass correlation coefficients ranging from 0.761 to 0.982 in sleep scoring with polysomnography (PSG) [22]. The montage was combined with four electroencephalogram derivations (Fp1–M2, Fp2–M1, Fp1– average M, and Fp2–average M), using Fp1–Fp2 and Fp2–Fp1 for left and right electrooculography, respectively, and M1–M2 for chin electromyography to analyze sleep staging. Sleep stages were scored blindly by experienced scorers according to the guidelines of the American Academy of Sleep Medicine (AASM) guidelines [23]. Sleep variables included sleep onset latency (SOL), total sleep time (TST), sleep efficiency (SE), wake after sleep onset (WASO), and the amount and proportion of each sleep stage, including non-rapid eye movement (NREM) sleep: stages 1 (N1), 2 (N2), and 3(N3), and REM sleep.

#### Body temperature and sweating rate

While acknowledging the non-uniformity of temperature on the tympanic membrane surface, the measurement of body temperature using the tympanic method (tympanic membrane temperature: TMT) represents a standard non-invasive method that properly approximates CBT [24,25]. The participants were instructed to wear a core body temperature sensor (BL100, Technonext Inc., Chiba, Japan) during night-time sleep. This sensor, with dimensions of 58 mm × 37 mm × 20 mm, is capable of measuring the TMT, providing an indication of deep-body temperature through an ear probe.

Furthermore, it continuously recorded sweating rates, skin surface temperatures, and skin humidity over time at a sampling frequency of 1 Hz during sleep and wirelessly transmitted the data to a computer via Bluetooth. The validity of this device for recording measurements during human sleep was confirmed [26].

As there were individual differences in TMT [27], skin temperature [28], and sweating rate [29], the values at bedtime at the start of the experiment were used as the baseline, and the subsequent measured values were analyzed for differences from the baseline. The variables recorded by the TMT sensor during sleep were computed on a per-min basis and then averaged within each 15-min interval. The values for each 15-min period were subsequently averaged across four time blocks: 11 pm–1 am, 1 am–3 am, 3 am–5 am, and 5 am–7 am. Furthermore, measurements obtained every 30 min after sleep onset were averaged for comparison.

#### Heart rate variability

Sympathetic nerve activity was estimated from the heart rate variability (HRV) of the electrocardiogram (ECG) data. For frequency analysis, the power within the low-frequency (LF; 0.04–0.15 Hz) and high-frequency (HF; 0.15–0.4 Hz) components of heart rate variability was obtained using Actiheart 5 monitors, (Cambridge Neurotechnology, Cambridge, UK). These wearable devices are flat and lightweight (10.5 g) and have demonstrated high levels of intra- and inter-instrument reliability, along with robust validity [30]. The participants were instructed to securely affix the Actiheart monitor to their chest using a belt during sleep at night. IBI data flagged as invalid by Actiheart Software ver.5; Cambridge Neurotechnology, Cambridge, UK) were excluded from the analysis.

Furthermore, the root mean square of successive differences (RMSSD) between adjacent R-R intervals was calculated. Vagal modulation is used to define the HF component, while the sympathetic and parasympathetic nervous systems modulate the LF components, indicating that sympathetic activity may be due to an increased LF component. The overall sympathovagal balance can be represented by the ratio between the LF and HF components, which determines the balance of the tone [31]. Similarly, in the evaluation of the CBT and perspiration rate, variables obtained from ECG recordings during sleep were computed on a per-minute basis and averaged within each 15-min interval. The values for each 15-min period were subsequently averaged across four time blocks: 11 pm–1 am, 1 am–3 am, 3 am–5 am, and 5 am–7 am. The values calculated every 30 min after sleep onset were averaged.

### Statistical analysis

Statistical analyses were performed using SPSS statistics version 29.0 (IBM SPSS Statistics for Windows; IBM Corp., Armonk, NY, USA). Data were normally distributed according to the Shapiro–Wilk test. For demographic variables, we calculated mean values and standard deviations.

The variables from the night before the experiment and the results of OSA-MA and EEG were compared between the two garment conditions and analyzed using a paired t-test. Moreover, the effect size statistic (d) was analyzed to determine the magnitude of the effect independent of the sample size [32]. Differences were interpreted using Cohen’s (d) guidelines as trivial (<0.2), small (0.2–0.6), moderate (0.6–1.2), large (1.2–2.0), very large (2.0–4.0), and huge (>4.0) [33].

The data recorded by the core body temperature sensor were analyzed and displayed in 15-min time bins, whereas the HRV data were analyzed and displayed in 2-hour time bins. For the time sequence data (TMT, perspiration, skin temperature, and HRV variables), the significance of the effects (garment type, time, and garment type × time) was analyzed using repeated measures ANOVA with a grouping factor (garment type). We adapted the values with the Greenhouse–Geisser correction if the Mauchly test was significant. Post-hoc pairwise comparisons were performed, and the results were interpreted using the Bonferroni correction.

## Results

### Pre-experiment and subjective sleep assessment

Total sleep time at the night prior the experiment was not statistically different between the two garment conditions ((FIR; 367.6 ± 104.39) min, control; (359.45 ± 71.27) min, *t*(14) = 0.208, *p* = 0.839).

Regarding the subjective sleep assessment evaluated by the OSA-MA, there were no significant differences between the FIR and control garments for all five factors (Table 1). However, when focusing on individual questions, the score of stress level (Question ‘I am stress free’ ‘I feel stressed’) showed better tendency with FIR compared than control (*t*(14)=2.120, *p*=0.052, d=0.936).

**Table 1.** Subjective sleep quality assessed by the OSA-MA version.

| Parameters | FIR | Control | t | p | d |
| --- | --- | --- | --- | --- | --- |
| Sleepiness on rising | 49.6 (6.6) | 48.4 (7.0) | 0.715 | 0.486 | 0.169 |
| Initiation and maintenance<br>of sleep | 39.5 (7.2) | 38.8 (12.8) | 0.207 | 0.839 | 0.066 |
| Frequent dreaming | 47.6 (9.5) | 50.6 (9.9) | 1.107 | 0.287 | 0.316 |
| Refreshing | 50.7 (6.7) | 47.7 (8.3) | 1.413 | 0.179 | 0.397 |
| Sleep length | 47.2 (8.1) | 45.4 (11.8) | 0.602 | 0.557 | 0.174 |

Abbreviations: FIR, Far-infrared; OSA, The Oguri Shirakawa and Azumi standard rating scale; MA, middle-aged and aged version.
Values are shown as the mean (standard deviation) for the FIR and control conditions.
Statistical significance was evaluated using paired t-test.

### Sleep Variables: Electroencephalogram analysis

Table 2 presents the sleep variables during the experimental night, as evaluated by EEG recording. %REM sleep (REM sleep percentage out of TST) was significantly larger in FIR compared with that in control (*t*(14) = 2.465, *p* = 0.027, d = 0.567). Sleep onset latency was shorter with FIR (33.1 ± 0.6 min) than with the control (43.7 min ± 0.7 min), though no statistical difference was observed. Except for % REM, sleep-related parameters for the overall 6 h of the EEG showed no significant differences between the two garment conditions.

**Table 2.** Comparisons of sleep-related variables between two garment conditions.

| Variables | FIR | Control | t | p | d |
| --- | --- | --- | --- | --- | --- |
| Total recording time<br>(TRT) (min) | 400.0 (6.7) | 402.6 (6.4) | 0.167 | 0.870 | 0.053 |
| Total sleep time (TST)<br>(min) | 366.7 (6.1) | 381.0 (6.4) | 0.708 | 0.490 | 0.253 |
| Sleep onset latency (min) | 33.1 (0.6) | 43.7 (0.7) | 0.981 | 0.343 | 0.255 |
| Sleep efficiency<br>(TST/TRT) (%) | 84.5 (12.2) | 84.7 (12.7) | 0.053 | 0.959 | 0.019 |
| Wake after sleep onset<br>(min) | 36.4 (0.6) | 27.7 (0.5) | 0.479 | 0.639 | 0.185 |
| N1 (min) | 31.7 (0.5) | 35.3 (0.6) | 0.715 | 0.486 | 0.230 |
| N2 (min) | 178.3 (3.0) | 201.0 (3.4) | 1.798 | 0.094 | 0.604 |
| N3 (min) | 73.1 (1.2) | 72.6 (1.2) | 0.041 | 0.968 | 0.013 |
| REM (min) | 83.3 (1.4) | 71.4 (1.2) | 1.620 | 0.128 | 0.379 |
| %N1 (%TRT) | 9.0 (5.0) | 9.4 (3.4) | 0.282 | 0.782 | 0.084 |
| %N2 (% TRT) | 48.6 (5.5) | 53.0 (8.1) | 1.631 | 0.125 | 0.636 |
| %N3 (% TRT) | 20.1 (8.4) | 18.91 (8.1) | 0.506 | 0.620 | 0.143 |
| %REM (% TRT) | 22.2 (6.5) | 18.6 (6.5) | 2.465 | 0.027 | 0.567 |

FIR, Far-infrared; N1, non-REM sleep stage 1; N2, non-REM sleep stage 2; N3, non-REM sleep stage 3; REM, rapid eye movement. Values are shown as mean (standard deviation) unless specified otherwise. Statistical significance was evaluated using paired t-test.

### Body temperature and sweating rate

Fig 1 shows the decrease from baseline for each variable across night sleep over time. The extent of the decrease in the TMT was greater in the FIR group than in the control group at all clock times. Although there was not a significant garment type x time interaction (*F*(26, 234)=0.411, *p*=0.996, *η*_*p*_^2^=0.044) in TMT, significant main effect in time (*F*(26, 234)=4.046, *p*<0.001,*η*_*p*_^2^=0.310), and tendency in type of garment (*F*(1, 9)=4.853, *p*=0.055, *`η`_p_*^2^=0.350), was observed (Fig 1A). The change in sweating rate from baseline showed a decreasing trend over time for the FIR group, whereas the control group showed an increasing trend. A significant garment x time interaction was found in sweating rate (*F*(26,572)=55.05, *p*<0.001, *η*_*p*_^2^=0.338). Additionally, significant main effect of sweating rate in time (*F*(26, 572)=4.853, *p*=0.050, *η*_*p*_^2^=0.714), in garment type (*F*(1, 22)=25.99, *p*<0.001, *η*_*p*_^2^=0.542), respectively (Fig 1B). Difference of skin temperature showed a significant main effect in time (*F*(26, 286)=4.046, *p*=0.013,*η*_*p*_^2^=0.139), though no significant interaction (Fig 1C). Skin humidity showed no significant garment × time interaction or main effect of the garment type (Fig 1D).

**Fig 1.**
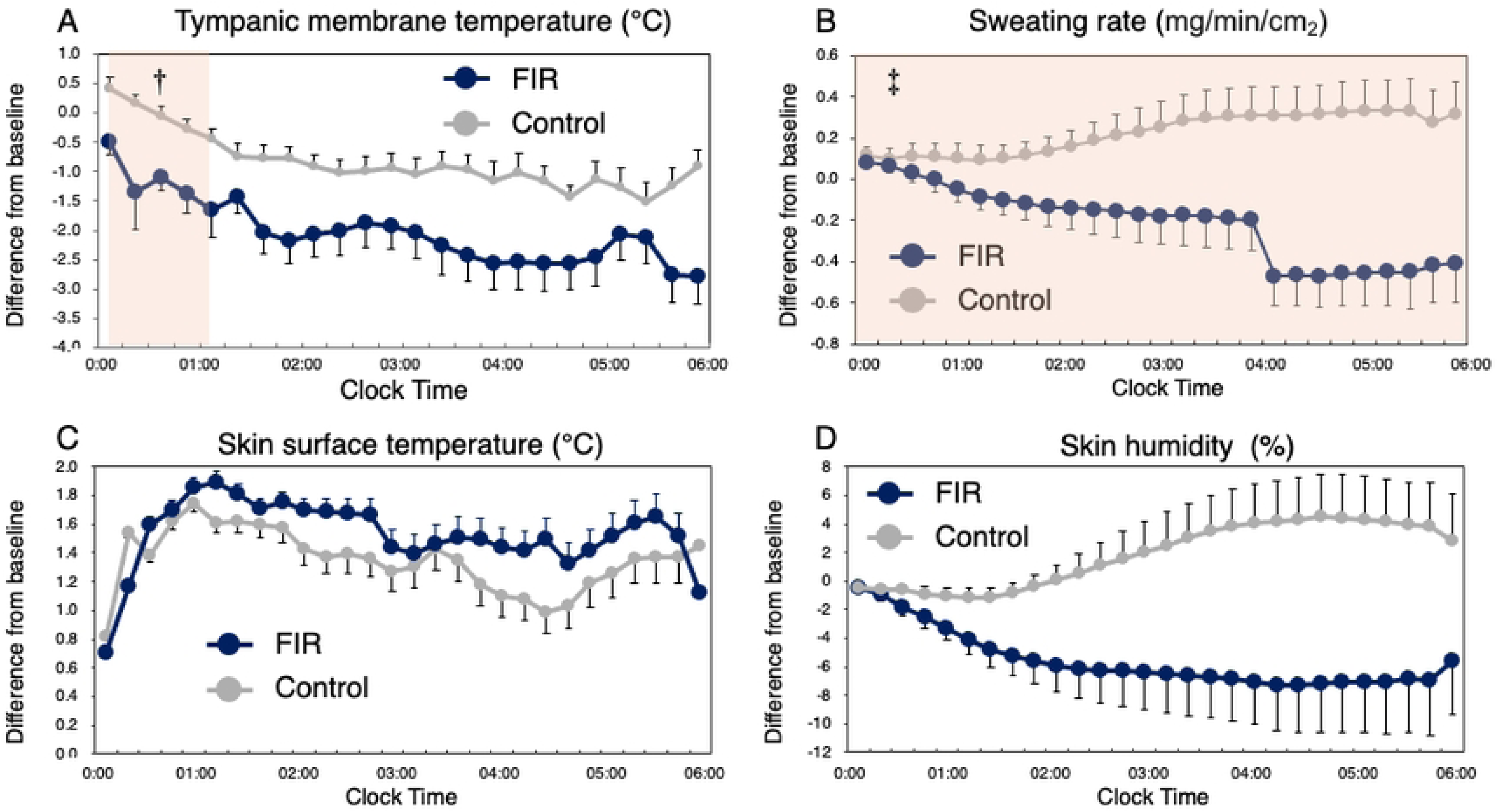
Comparisons of difference from baseline in measured variables over actual time clock. (A) Tympanic membrane temperature (TMT), (B) sweating rate, (C) skin surface temperature, (D) skin humidity. Two-way repeated measures of ANOVA with Bonferroni’s multiple comparison test showed a significant main effect of time in TMT and a garment type x time interaction in sweating rate. Blue circle and line indicate FIR and gray control, respectively. Error bar indicates standard error. † P<0.05, ‡ P<0.01 versus control (Bonferroni adjusted t-test).

Concurrently, as individual differences in the latency to sleep onset were observed, changes in the variables from sleep onset over time are shown in Figure 2. The trend of the results is similar to the change over time, change of TMT showed a significant main effect in time (*F*(4.23, 80.45)=4.616, *p*=0.002,*η*_*p*_^2^=0.195), in garment type (*F*(1, 19)=10.85, *p*=0.004,*η*_*p*_^2^=0.363), though no significant garment x time interaction was observed (*F*(4.23, 80.45)=0.370, *p*=0.839, *η*_*p*_^2^=0.019) (Fig 2A). Change of Perspiration rate revealed a significant garment type x time interaction (*F*(1.71, 32.4)=0.901, *p*=0.026,*η*_*p*_^2^=0.186), main effect in garment type (*F*(1, 19)=5.83, *p*=0.026,*η*_*p*_^2^=0.235), respectively (Fig 2B). No significant interaction or main effect was observed on the change in skin temperature (Figure 2C). Regarding skin humidity, there was a main effect in garment type (*F*(1, 20)=4.68, *p*=0.043,*η*_*p*_^2^=0.190) (Fig 2D).

**Fig 2.**
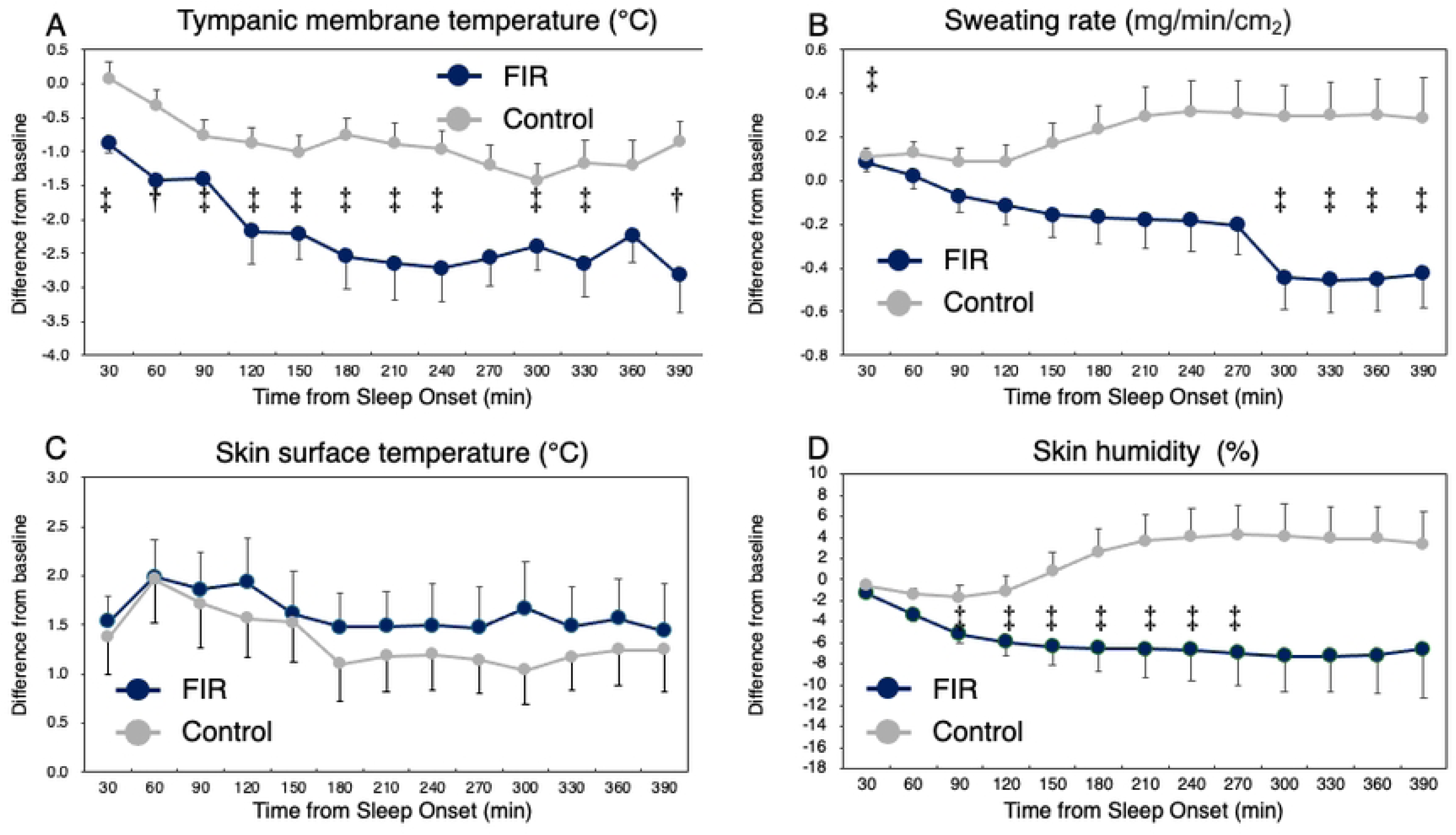
Comparisons of difference from baseline in measured variables after time of sleep onset. (A) Tympanic membrane temperature (TMT), (B) Sweating rate, (C) Skin surface temperature, (D) Skin humidity. Two-way repeated measures of ANOVA with Bonferroni’s multiple comparison test showed a significant main effect of time in TMT and a garment type x time interaction in sweating rate. Blue circle and line indicate FIR and gray control, respectively. Error bar indicates standard error. † P<0.05, ‡ P<0.01 versus control (Bonferroni adjusted t-test).

### Heart rate variability

Fig 3 shows the HRV variables averaged over the four time blocks during sleep. Although RMSSD manifested a significant main effect in time (*F*(1.52, 39.50)=4.24, *p*=0.027,*η*_*p*_^2^=0.145), there was no significant garments type x time interaction (*F*(1.52, 39.50)=1.22, *p*=0.296,*η*_*p*_^2^=0.045) nor main effect in garments type (*F*(1, 26)=0.003, *p*=0.027,*η*_*p*_^2^=0.00). Regarding LF, a significant garment type x time interaction (*F*(1.87, 48.57)=3.36, *p*=0.046,*η*_*p*_^2^=0.114). Post-hoc analysis revealed that the LF value of FIR was higher at 23:00-1:00 than that of control (p=0.027). LF/HF showed a significant main effect in time (*F*(1.44, 37.44)=4.43, *p*=0.029,*η*_*p*_^2^=0.146), there was no significant garments type x time interaction (*F*(1.44, 37.44)=1.08, *p*=0.332,*η*_*p*_^2^=0.040) nor main effect in garments type (*F*(1, 26)=0.026, *p*=0.872,*η*_*p*_^2^=0.001). HF showed no significant interaction (*F*(1.80, 46.74)=1.34, *p*=0.270,*`η`_p_*^2^=0.049) nor main effect across the time block (time; (*F*(1.80, 46.74)=1.35, *p*=0.268,*η*_*p*_^2^=0.049, garment, *F*(1, 26)=0.002, *p*=0.964,*η*_*p*_^2^=0.000).

**Fig 3.**
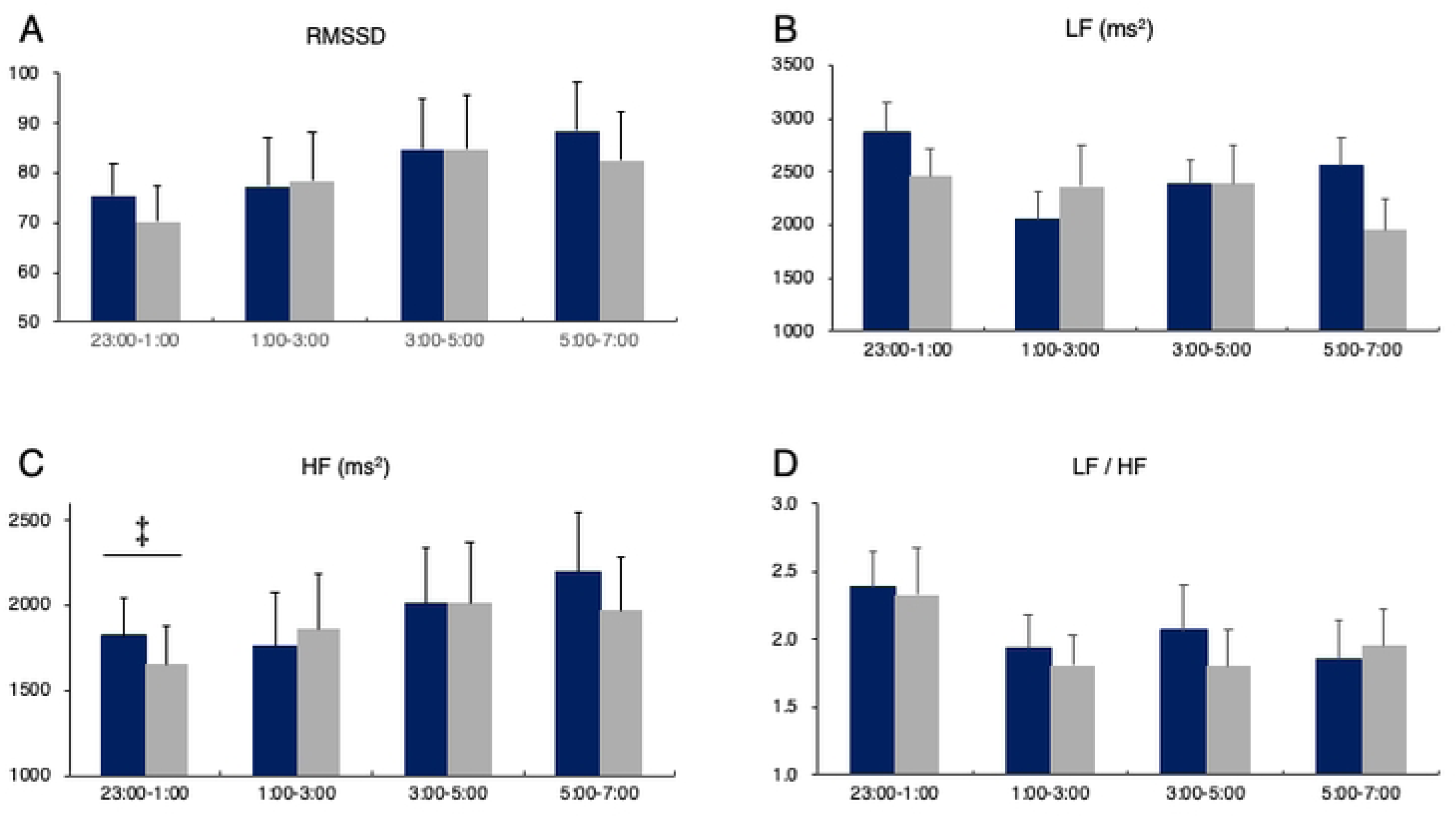
Change in heart rate variability variables across for time blocks for each garment condition. Blue histogram indicates FIR and gray histogram control, respectively. Error bar indicates standard error. RMSSD, root mean square of successive differences; LF, low-frequency; HF, high-frequency. † P<0.05 versus control (Bonferroni adjusted t-test).

## Discussion

The present study evaluated the effect of an FIR-emitting garment on physiological indices during night sleep relative to those observed during sleep in polyester clothes in healthy young populations. The proportion of REM sleep was significantly higher in the FIR group than in the control group. FIR provided significantly lower TMT throughout night sleep, a lower sweating rate in the final phase of sleep, and lower skin humidity in the middle of sleep compared to the control. Additionally, heart rate variability analysis revealed a significantly higher LF component in the initial phase of sleep compared to the control condition, suggesting that FIR may stimulate the sympathetic and parasympathetic nervous systems during the first cycle of NREM sleep.

The trends in TMT, sweating, skin temperature, and skin humidity during sleep were similar, although the plots shown in terms of time and elapsed time from sleep onset were slightly different. A previous study comparing PSG data from a young population showed robust commonalities in sleep-related variables other than slow-wave sleep [34]. In the current study, owing to the longer latency to sleep onset in the control group than in the FIR group, although there were no significant differences, assessing the time elapsed since sleep onset would fit the evaluation of the physiological trend. Ample evidence demonstrates the functional importance of the first NREM sleep cycle during sleep [35]. Sleep periods occur during the circadian phase of reduced CBT, in coordination with the circadian clock regulated by the hypothalamic suprachiasmatic nucleus [36]. CBT normally decreases, while it tends to increase in collaboration with the physiological processes of the body surface that activate the circulation of blood vessels and evaporation, resulting in a decrease in CBT [37]. In the present study, TMT, which corresponds to CBT, remained relatively low from the beginning to the end of sleep, suggesting that FIR ameliorate sleep propensity by incorporating proper thermoregulation. Although FIR have been shown to increase skin surface temperature [38], the effect of FIR on CBT remains unclear. Considering that FIR would improve local microcirculation [39], the observed decrease in TMT during sleep may be due to heat dissipation involved in facilitated skin vessel circulation.

The FIR-emitting garment accounted for a relatively larger proportion of REM sleep, which occurs more frequently in the latter half of night sleep by homeostatic regulation, as well as by the circadian clock [40], compared to the control. Generally, the CBT decreases and tends to persist at lower levels until it reaches the point of awakening [41]. Animal study demonstrated that REM sleep incremented brain temperature resulting from the relative increase in warmer vertebral, over carotid artery, blood flow [42], and accompanied by an increase in brain temperature of approximately 0.1–0.2°C [43]. In the FIR condition, body temperature was maintained at a lower level during REM sleep despite low levels of sweating that allowed the dissipation of inner body heat, apparently coordinating it as a suitable environment for restorative sleep. Sweating is the most effective autonomic thermoregulatory factor that provides the potential for heat loss via evaporation [44]. Thermoregulatory sweating during night sleep occurs substantially because of cutaneous vasodilation, causing increased blood flow to the body surface [45].

The sensitivity of the thermoregulatory system during REM sleep has been proven to be lower than that during slow-wave sleep, presumably due to an increase in the hypothalamic set-point temperature [46] and blunted sweating response during REM sleep [47]. A previous study reported that the amount of skin temperature change in humans was smaller during REM sleep than during NREM sleep, resulting in inadequate thermoregulation for maintaining sleep [48]. Therefore, the use of FIR garments may help maintain lower core body temperatures despite a relative increase in REM sleep, which disrupts sweating and thermoregulation. It is also essential to maintain adequate REM sleep, as reduced REM sleep has been linked to poor mental and physical health outcomes [49,50], increased risk of dementia [51], and all-cause mortality [52]. Taken together, even during prolonged REM sleep, in which the thermoregulatory mechanisms are placed in an unstable state, FIRs may have potential mechanisms to maintain a low core body temperature without resorting to sweating, which may involve the dilation of surface microvessels.

HRV analysis revealed that the LF components were higher in the FIR-emitting garment group than in the control group, specifically during the initial phase of sleep. LF components reflect the combined activity of the sympathetic and parasympathetic nervous system [31]. Although few studies have examined the association between FIR and the autonomic nervous system, Peng et al. explored the effect of FIR on foot skin surface temperature and heart rate variability, demonstrating that LF and HF activity significantly increased in the FIR group but not in the LF/HF ratio [38]. The current results indicate that FIR can increase autonomic nervous system (ANS) activity and maintain equilibrium. Previous studies have demonstrated that the FIR increases peripheral blood perfusion and maintains the LF/HF ratio in equilibrium [53]. The nonthermal effect of FIRs induces an increased NO2 concentration in the blood, which promotes epidermal vasodilation [54]. Moreover, FIR therapy induces significant HRV responses corresponding to increased LF, which is in agreement with our results. Given that HF and RMSSD, which reflect parasympathetic activity, did not differ, FIR yielded sympathetic dominance in the early stages of sleep, which was shown to decrease sympathetic tone [55]. However, because the LF/HF ratio is in equilibrium, it is possible that even the benefit from thermoregulation by dilated microvessels due to FIR emission may surpass increased sympathetic activity, eventually facilitating restorative sleep.

The current study has some limitations. First, the participants were limited to male participants. Women are concerned about menstrual cycles and cold extremities due to poor peripheral circulation, which is an issue in controlling sleep experiments. Therefore, a specific study on garments focused on female is warranted. Second, analyses of CBT, sweating, and HRV were not performed using the data for the epochs of sleep stages. In this study, physiological indices were analyzed based on elapsed sleep time owing to technical issues. Nevertheless, the present results demonstrate the efficacy of FIR-emitting sleepwear, as there is a trend toward a predominant sleep stage according to the sleep course. Third, although clothing of a similar color, shape, and texture was used as the control garment, concerns remained as to whether they were completely blind to the participants. However, no remarkable difference was observed in the subjective evaluation, which suggests that the placebo effect was avoided owing to the recognition of the FIR condition prior to the experiment. Regardless of these limitations, our results suggest that FIR-emitting sleepwear has beneficial effects on sleep and its associated physiology. Future studies should explore the effects of FIR-emitting sleepwear on sleep quality, encompassing populations ranging from general to specific purposes, such as those who are not satisfied with sleep, bedclothes in hospitals, and recovery garments for athletes in the sports field.

## Conclusion

The FIR-emitting garment serves to reduce TMT not only at sleep onset but also at a lower value throughout sleep, which is favorable for physiological sleep optimization. The maintenance of a lower TMT, despite low sweating, may be due to appropriately implemented thermoregulation during REM sleep, which promotes epidermal vasodilation. The environment under wearing FIR-emitting garments has the potential to facilitate restorative sleep, despite increased sympathetic activity, which may be offset by optimizing the peripheral circulatory environment using FIR. These physiological alterations provided by the suitable sleeping environment may overall promote the general health related to well-being.

## Acknowledgements

The authors are grateful to Atsushi Ichinose for his help with the experiment and to Takashi Sakata for technical cooperation.

## Author Contribution

MN: Research organization, project design, and result description; MN,TN, SS, and SY: Data acquisition, data interpretation; MN: Research methodology, statistical analysis, and result description. MO and KF prepared the manuscript All the authors contributed to the manuscript and approved the submitted version.

